# Hippo pathway-mediated YAP1/TAZ inhibition is essential for proper pancreatic endocrine specification and differentiation

**DOI:** 10.1101/2022.05.31.494216

**Authors:** Yifan Wu, Kunhua Qin, Kevin Lopez, Jun Liu, Michael Nipper, Janice J Deng, Xue Yin, Logan Ramjit, Zhengqing Ye, Pei Wang

## Abstract

The Hippo pathway plays a central role in tissue development and homeostasis. However, the function of Hippo in pancreatic endocrine development remains obscure. Here, we examined the roles of Hippo pathway mediated YAP1/TAZ inhibition in the development stages of endocrine specification and differentiation. While YAP1 protein was localized to the nuclei in bipotent progenitor cells, Ngn3 expressing endocrine progenitors completely lost YAP1. Using genetic mouse models, we found that this inactivation of YAP1 requires both intact Hippo pathway and NGN3 protein. Deleting the Lats1 and 2 kinases (*Lats1&2*) in endocrine progenitor cells of developing mouse pancreas with Ngn3-Cre blocked endocrine progenitor cell differentiation and specification, resulting in reduced islets size and disorganized pancreas at birth. Loss of Lats1&2 in NGN3 expressing cells activated YAP1/TAZ transcriptional activity and recruited macrophages to the developing pancreas. These defects were rescued by deletion of YAP1/TAZ genes, suggesting that tight regulation of YAP1/TAZ by Hippo signaling is crucial for pancreatic endocrine specification. In contrast, deletion of Lats1&2 using beta-cell specific *MIP1*^*CreER*^ resulted in a phenotypically normal pancreas, indicating that Lasts1&2 are dispensable for pancreatic beta-cell function. Our results demonstrate that YAP1/TAZ inhibition in the pancreatic endocrine compartment is not a passive consequence of endocrine specification. Rather, Hippo pathway-mediated YAP1/TAZ inhibition in endocrine progenitors is a prerequisite for endocrine specification and differentiation.

## Introduction

The mammalian pancreas is composed of exocrine and endocrine compartments, which both originate from embryonic endoderm. The exocrine pancreas is mainly comprised of acinar cells, which secrete various digestive enzymes, and ductal cells which transport these enzymes to the duodenum. The endocrine pancreas, which consists of α-, β-, δ-, PP- and ε-cells, produces several hormones that are secreted into the blood with organism-wide functions. Of these functions, regulation of glucose homeostasis is predominately associated with pancreatic endocrine function. Developmental defects in the pancreas lead to devastating pathological diseases including congenital pancreas abnormalities, congenital hyperinsulinism, and neonatal diabetes[1–3].

Initially discovered in Drosophila, the Hippo signaling pathway is best known for its roles in suppressing growth, promoting apoptosis, and regulating organ size[4]. In mammals, this pathway has more complicated, context-dependent functions with regard to regulating tissue homeostasis[5]. The mammalian Hippo pathway is comprised of a kinase signaling cascade beginning with the Ste-20-like protein kinases (MST1&2), which directly phosphorylate and activate Large tumor suppressors 1&2 (LATS1&2)[5]. Upon activation, LATS1&2, the final kinases in this cascade, directly phosphorylate the effectors of the Hippo pathway including the transcription coactivators Yes-Associated Protein 1 (YAP1) and WW-domain-containing transcription regulator 1 (WWTR1, also known as TAZ). The phosphorylation of YAP1 and TAZ facilitates these effectors to undergo proteasome-mediated degradation or cytosolic sequestration in opposition to nuclear translocation[6]. In contrast, inhibiting the Hippo pathway stabilizes YAP1 and TAZ, thereby enabling their cytosol-to-nucleus translocation and initiating the downstream transcriptional program. This transcriptional activation involves physical interaction of other transcription factors with YAP1/TAZ such as TEADs [5]. Prior studies have shown that compared with inactivation of upstream components of the Hippo pathway such as *Mst1&2*, specific genetic inactivation of *Lats1&2* more robustly facilitates YAP1 and TAZ transcriptional activities[6].

The Hippo signaling pathway has been shown to play key roles in pancreatic development, both in cell lineage differentiation and pancreatic morphogenesis[7,8]. During early-stage pancreatic development, deletion of *Mst1/2* in pancreatic progenitor cells results in dysregulation of acinar expansion. De-differentiation of acinar cells into ductal-like cells (also known as acinar-to-ductal metaplasia, or ADM), immune cell infiltration, and pancreatic auto-digestion have also been observed following *Mst1/2* deletion. However, the overall function of the islets is not affected by the loss of *Mst1/2* [9,10]. *Lats1&2* also serve vital functions in early pancreatic development before the secondary transition when endocrine differentiation reaches the peak [11]. Endocrine specification is driven by the basic helix-loop-helix transcription factor Neurogenin 3 (Ngn3) in bipotent pancreatic progenitors at secondary transition stage [12,13]. Ngn3 is a master regulator of pancreatic islet differentiation and initiates stepwise cell fate determination, delamination, and migration of differentiating endocrine cells [14–16]. *In vitro* culture experiments found that NGN3 directly represses the transcription of YAP1[17]. These observations lead to the notion that Hippo pathway-mediated YAP1 inhibition in NGN3+ endocrine progenitors is dispensable for the subsequent endocrine differentiation process, but this has not been examined with *in vivo* developmental models [7].

Here, we investigated the function of Hippo in endocrine cell development and homeostasis by inactivating *Lats1&2* in NGN3+ endocrine progenitor cells and β-cells. We found that Hippo activity in NGN3+ cells is required for mouse endocrine progenitor specification and differentiation, but is not necessary for pancreatic β-cell function. By further introducing *Yap1/Taz* deletion into *Lats1&2* null cells, we demonstrated that attenuation of YAP1/TAZ through genetic ablation rescued the observed phenotypes driven by *Lats1&2* inactivation. Our results uncovered a context-dependent function of the Hippo signaling pathway in maintaining normal development and tissue homeostasis in the pancreas.

## Materials and methods

### Ethics Statement

All procedures have been approved by the Institutional Animal Care and Use Committee (IACUC) at The University of Texas Health San Antonio.

### Generation of conditional knockout mice

All animal study protocols were approved by the UT Health San Antonio Animal Care and Use Committee. *Ngn3*^*Cre*^ mice (stock number: 005667), *MIP*^*CreER*^ (stock number: 024709), and *R26R*^*EYFP*^ mice (stock number: 006148) were obtained from The Jackson Laboratory. *Ngn3*^*Cre*^ mice were kindly provided by Dr. Andrew Leiter and Dr. Seung Kim. *Lats1*^*fl/fl*^ and *Lats2*^*fl/fl*^ mice were kindly provided by Dr. Randy L. Johnson. *YAP1*^*fl/fl*^ and *Taz*^*fl/fl*^ mice were kindly provided by Dr. Eric N. Olson. We generated (1) *Ngn3*^*Cre*^*Rosa26*^*LSL-YFP*^*Lats1*^*fl/fl*^*Lats2*^*fl/+*^ mice (as Control), (2) *Ngn3*^*Cre*^*Rosa26*^*LSL-YFP*^*Lats1*^*fl/fl*^*Lats2*^*fl/fl*^ *mice* (NL mice), (3) *Ngn3*^*Cre*^*Rosa26*^*LSL-YFP*^*Lats1*^*fl/fl*^*Lats2*^*fl/+*^*YAP1*^*fl/fl*^*Taz*^*fl/fl*^ mice (NTY mice), and (4) *Ngn3*^*Cre*^*Rosa26*^*LSL-YFP*^*Lats1*^*fl/fl*^*Lats2*^*fl/fl*^*YAP1*^*fl/+*^*Taz*^*fl/fl*^ mice (NLT mice). (5) *Ngn3*^*Cre*^*Rosa26*^*LSL-*^ Y^*FP*^*Lats1*^*fl/fl*^*Lats2*^*fl/fl*^*YAP1*^*fl/fl*^*Taz*^*fl/fl*^ mice (NLTY mice). (6) *MIP*^*CreER*^*Rosa26*^*LSL-YFP*^*Lats1*^*fl/fl*^*Lats2*^*fl/fl*^ *mice* (ML mice). All offspring were genotyped by PCR of genomic DNA from the toe with primers specific for the *Ngn3*^*Cre*^, *MIP*^*CreER*^, *Rosa26*^*LSL-YFP*^, *Lats1, Lats2, Yap1*, and *Taz* loci. For the timed mating experiment, male mice were introduced to the cage in the afternoon and removed in the morning on the second day, which was considered gestational day 0.5 (E0.5). The pregnant mice were then used to harvest the embryos at the indicated time. For the tamoxifen-mediated *Lats1&2* deletion in adult β-cells, 6-8 weeks old mice were intraperitoneal (i.p) injected with 200 mg/kg tamoxifen (TAM, Sigma-Aldrich, T5648) for 5 consecutive days. For the early β-cell *Lats1&2* deletion, female *Lats1&2*^*fl/fl*^ mice and male mice carrying *MIP*^*CreER*^; *Lats1&2*^*fl/fl*^ were timed mated and after 12 days, the pregnant mice were injected with 100 mg/kg tamoxifen once[18]. PCR was used for validation of knockout alleles (S Table 2).

### H&E staining, Picrosirius red staining, immunofluorescence, and immunohistochemistry

Pancreases harvested from animals were fixed in 4% paraformaldehyde overnight and then submitted to the Histology Core of the University of Texas Health Science center at San Antonio. The tissues were immersed in serial dilutions of ethanol and xylene and then embedded in the paraffin. The blocks were then cut into 5 µm sections with a microtome. H&E staining was performed at the Histology Core. For Picrosirius red staining, the section slides were first deparaffinized with xylene and washed in serial dilutions of ethanol solutions. Then, the slides were stained with Hematoxylin, 0.1% Fast Green, and Pico-Sirius Red solution (Polysciences, 24901) as previously described[19].

For Immunofluorescent staining, tissues were deparaffinized, rehydrated, and submerged in 200°C heated R-Universal Epitope Recovery Buffer solution (Electron Microscopy Sciences, Hatfield, PA, 62719-20) for 30 minutes and then let cool at room temperature for 25 minutes. Sections were permeabilized using 0.3% TBST (0.3% Triton X-100, Acros Organics, Fair Lawn, NJ, 21568-2500) and 0.025% PBST (0.025% Tween20, Fisher Bioreagents, Fair Lawn, NJ, BP337-500) for 4 minutes each. Sections were subsequently blocked with 10% donkey serum in 0.025% PBST for 35 minutes at room temperature. Sections were then incubated with primary antibodies diluted in 10% donkey serum in 1X PBS at 4°C overnight. Then, sections were incubated with fluorescent-tagged Alexa Fluor secondary antibodies (1:250, Jackson ImmunoResearch, West Grove, PA) diluted in 10% donkey serum in 1X PBS for 1 hour at room temperature. Additionally, sections were incubated with DAPI (1:1000, Invitrogen, Carlsbad, CA, P36935) for 4 minutes at room temperature. Finally, sections were covered with a drop of VectaShield Vibrance Antifade Mounting Medium (Vector Laboratories, Inc., Burlingame, CA, H-1700). All images were captured using Microsystems DMI6000 B microscope and software (Leica Microsystems, Buffulo Grove, IL) and Zeiss LSM510 confocal microscopes. All primary and secondary antibodies used are listed in Table 1.

**Table 1:**

| **Primary Antibody** | **Company** | **Catalog #** | **Dilution** | **Application** |
| --- | --- | --- | --- | --- |
| alpha-Amylase (AMY) | Sigma Aldrich | A8273 | 1:500 | IF |
| alpha-SMA (ACTA2) | Santa Cruz Biotechnology | 53142 | 1:250 | IF |
| beta-Catenin (CTNNB1) | BD Biosciences | 610153 | 1:100 | IF |
| CD45 | Biolegend | 103102 | 1:50 | IF |
| KRT19 | DSHB | Troma-III | 1:50 | IF |
| E-Cadherin (CDH1) | Cell Signaling Technology | 3195S | 1:500 | IF |
| F4/80 | Cell Signaling Technology | 70076S | 1:50 | IF |
| GFP | Aves | GFP-1020 | 1:500 | IF |
| Insulin (INS) | Abcam | ab7842 | 1:250 | IF |
| ISL1/2 | DSHB | 39.4D5 | 1:50 | IF |
| Ki67 | Biolegend | 652401 | 1:25 | IF |
| NGN3 | DSHB | F25A1B3 | 1:50 | IF |
| PDX1 | DSHB | F109-D12 | 1:50 | IF |
| Somatostatin (SST) | Santa Cruz Biotechnology | sc-7819 | 1:100 | IF |
| SOX9 | Cell Signaling Technology | 82630S | 1:50 | IF |
| SPP1 | RD Systems | AF808 | 1:50 | IF |
| TAZ | Sigma Aldrich | HPA007415-100UL | 1:50 | IF |
| YAP | Cell Signaling Technology | 14074S | 1:100 | IF |

### RNA extraction and Reverse transcription real-time PCR analysis (RT-qPCR)

Pancreases from the animal were homogenized in Trizol (Invitrogen, 15596026) using a probe sonicator from Qsonica (25%, 15s ON, 15s OFF) immediately after harvest. RNA extraction and RT-qPCR were performed as previously described[20]. The primers for quantitative real-time PCR were listed in the supplemental table S1

### Western blot analysis

Islets from *Lats1&2*^*fl/fl*^ and *MIP1*^*CreER*^; *Lats1&2*^*fl/fl*^ mice were isolated as described[21]. After three hours of recovery, 50-60 islets were handpicked into low retention tubes (Eppendorf, Z666548). Total protein was extracted by incubating the islets with Laemmli buffer at 95°C for 5 mins and homogenized for another 5 mins in a sonication device (Diagenode, B01060001) at 4°C. Protein concentration was determined by the BCA method (Thermo Fischer Scientific, 23225) and 5-10 µg of total protein were loaded onto the 10% SDS-PAGE gel for electrophoresis. (All antibody information and working concentrations are shown in S1 Table).

### Blood glucose level and glucose tolerance test

Blood glucose levels of the animal were determined by a glucometer (BIONIME GS550). The glucose tolerance test was performed on overnight fasted 5-month post-tamoxifen *Lats1&2*^*fl/fl*^ and *MIP1*^*CreER*^; *Lats1&2*^*fl/fl*^ mice. 1.5 g/kg glucose was Intraperitoneal injected into animals and blood glucose levels were monitored at indicated time points in the figures.

### Statistical analysis

All results in this study were presented as the mean ± standard error of the mean (s. e.m.). Statistical analysis was performed by a two-tailed Student’s t-test. p values < 0.05 were considered statistically significant.

### Additional materials

Detailed reagents information can be found in the S1 Text.

## Results

### Inactivation of *Lats1&2* in NGN3+ cells led to defects in mouse postnatal growth, acinar atrophy, and ductal expansion in the pancreas

To determine the roles of *Lats1* and *Lats2* in endocrine progenitor cells and pancreas development, both genes were inactivated in the *Ngn3*^*Cre*^ transgenic mouse strain. For tracing of Cre-recombinase activity, we also crossed the *Rosa26*^*LSL-YFP*^ line. The heterozygote mice (*Ngn3*^*Cre*^*Lats1*^*fl/+*^*Lats2*^*fl/+*^*Rosa26*^*LSL-YFP*^) and single knockout mice (*Ngn3*^*Cre*^*Lats1*^*fl/fl*^*Lats2*^*fl/+*^*Rosa26*^*LSL-YFP*^ or *Ngn3*^*Cre*^*Lats1*^*fl/+*^*Lats2*^*fl/fl*^*Rosa26*^*LSL-YFP*^) from the initial F1 and F2 generations showed no abnormal phenotype. We, therefore, bred the single knockout mice with the *Lats1&2* conditional allele line to get the *Ngn3*^*Cre*^*Lats1*^*fl/fl*^*Lats2*^*fl/fl*^*Rosa26*^*LSL-YFP*^ mice (abbreviated as NL mice). The *Lats1* single knockout littermates were used as control (Ctrl) (Figure S1A). NL mice were born in Mendelian ratios with no obvious differences from the littermate control. However, starting from one week of age, it became obvious that the NL mice were smaller in size than the control mice. By the time of postnatal day 19 (P19), the NL mice appeared smaller and weaker than their littermates (Figure S1B), with a much smaller pancreas (Figure S1C), and suffered from hypoglycemia. The majority of NL mice died before three weeks of age, exhibiting rear limb weakness or paralysis. Feeding the mice with 10% glucose[24] or keeping them with their parents delayed, but did not prevent their death (data not shown), indicating that early death was only partially due to hypoglycemia. A few mice survived beyond three weeks with much smaller body sizes and infertility. These growth-related phenotypes could be partially attributed to the expression of *Ngn3*^*Cre*^ in the central nervous system[25].

Histological analysis with H&E staining showed that the acinar cells, ductal system, and islets could be clearly identified in control pancreases, whereas remarkably disorganized structures were seen in the center of the NL pancreases (Figure 1A) including noticeable expansion of ductal cells and fewer acinar cells (Figure S1E). Immunohistochemical staining of Insulin (INS) revealed much smaller islets in NL pancreases in addition to structural disorganization, evident by immunofluorescent co-staining of Insulin and Glucagon (GCG) (Figure 1B). We further analyzed the expression level of lineage-specific genes in the P1 pancreas using quantitative RT-PCR (qPCR). We found that the expression of acinar-specific markers, *Ptf1a*, Amylase (*Amy*), and carboxypeptidase A1 (*Cpa1*), as well as endocrine-specific markers, Chromogranin A (*ChrA*), Insulin 1 (*Ins1*), and Insulin 2 (*Ins2*) were significantly lower in NL mice when compared with control mice (Figure 1C). In comparison, while the expression of two ductal-specific markers, *Hnf1b* and *Sox9*, had no significant change, *Krt19* was significantly higher in NL pancreases (Figure 1C). To pinpoint the starting time of the observed defect in NL mice, we examined the E15.5 pancreas, when expression of Ngn3 reaches its peak. We did not observe any obvious differences between NL and control pancreases at E15.5 by H&E staining. The expansion of central ductal structure was noted in the NL pancreas at E16.5 which became even more obvious at P1 from H&E staining and KRT19 immunohistochemistry staining (Figure 1D). Together, these observations suggest that genetic disruption of *Lats1&2* in NGN3+ cells leads to acinar atrophy and ductal expansion in the postnatal pancreas, which are likely initiated in the developing embryonic pancreas.

**Figure 1.**
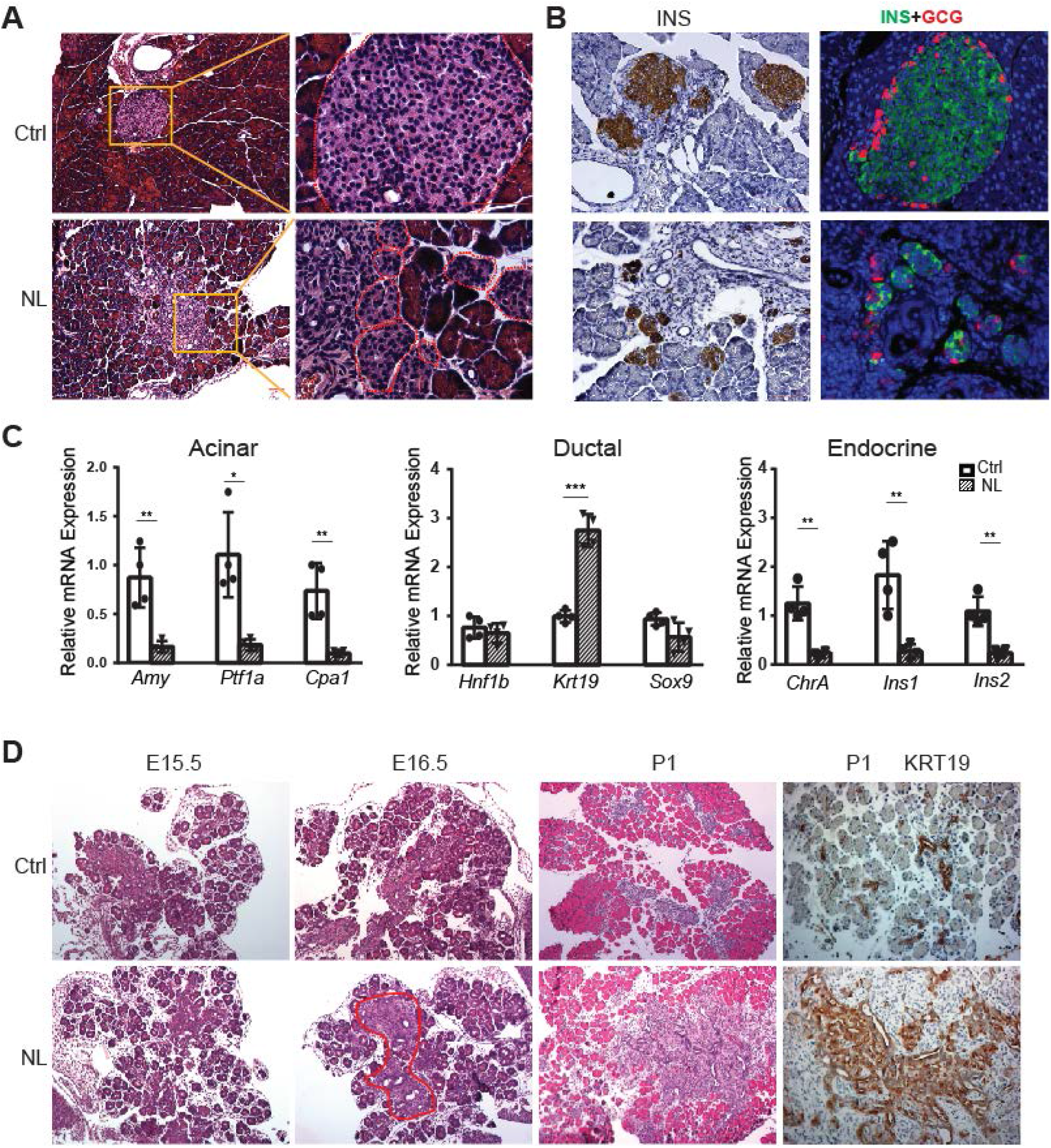
Deletion of *Lats1/2* by *Ngn3*^*Cre*^ perturbs pancreatic endocrine development. A. H&E staining of pancreatic tissue from NL mice showed phenotypically abnormal pancreatic islets as compared to those in control mice. B. Immunostaining of Insulin (INS) and Glucagon (GCG) showed reduced size of pancreatic islets and decreased abundance of β-cells in NL mice. C. Quantitative RT-PCR analysis of the expression levels of acinar, ductal, and endocrine genes in neonatal (P1) control and NL pancreas (n=4). * p<0.05; ** p<0.01; *** p<0.001. D. H&E staining and immunohistochemical staining of KRT19 of pancreatic tissue from control and NL mice at indicated stage. Structural changes in histology and a higher level of KRT19 in NL mice were noted at the indicated stages. Scale bar: 100 μm.

### Inactivation of *Lats1&2* blocks endocrine specification and differentiation

In the developing pancreas, expression of *Ngn3* marks the beginning of endocrine cell differentiation and reaches its peak at E15.5. However, as described before, apparent defects in pancreatic morphology were not observed through H&E staining until E16.5. Therefore, subsequent analysis was focused on the E16.5 pancreas. We monitored Cre-recombinase activity with *Rosa26*^*LSL-YFP*^ allele. YFP expression was used as an indicator for *Lats1&2* deletion as no viable antibodies were available for staining of LATS1&2. We also performed immunofluorescent staining of pancreatic and duodenal homeobox 1 (PDX1), an early pancreatic progenitor marker and β-cell-specific transcriptional factor. YFP+ cells formed cell clusters and were co-stained with PDX1 in control pancreases (Figure 2A). In contrast, in NL pancreases, the majority of YFP+ cells did not form clusters, but instead formed a distinct layer next to ductal cells. In addition, there were also much fewer PDX1 and YFP double-positive cells in NL pancreases, as compared with the control (Figure 2A), suggesting that most endocrine progenitors in the NL pancreas do not develop into β-cells at E16.5. Presence of a few double-positive cells in NL pancreases could be due to incomplete deletion of genes. At birth, control pancreases showed co-staining of YFP and PDX1 in the center region and YFP single positive cells in the periphery region, which is consistent with the normal structure of Islet of Langerhans where β-cells localize to the center of islets while α-cells localize to the periphery. By contrast, NL pancreases had much fewer PDX1 positive cells which were found in small unorganized cell clusters, while the majority of YFP+ cells did not express PDX1 (Figure S2A). To further delineate the defect of *Lats1/2* null endocrine progenitor cells, we performed immunostaining on the transcription factors Islet 1 (ISL1) and NK2 homeobox 2 (NKX2.2), which are expressed after NGN3 in endocrine pancreas development [26]. We found that ISL1 was widely expressed in YFP+ cells in control pancreases, but only expressed in a few small YFP+ cell clusters in NL pancreases, in which the majority of YFP+ cells did not express ISL1 (Figure 2B). A similar expression pattern was also observed for NKX2.2 (Figure 2C). Thus, these data indicate that loss of *Lats1&2* leads NGN3+ endocrine progenitor cells to halt the differentiation program almost immediately and prevents them from further endocrine lineage progression.

**Figure 2.**
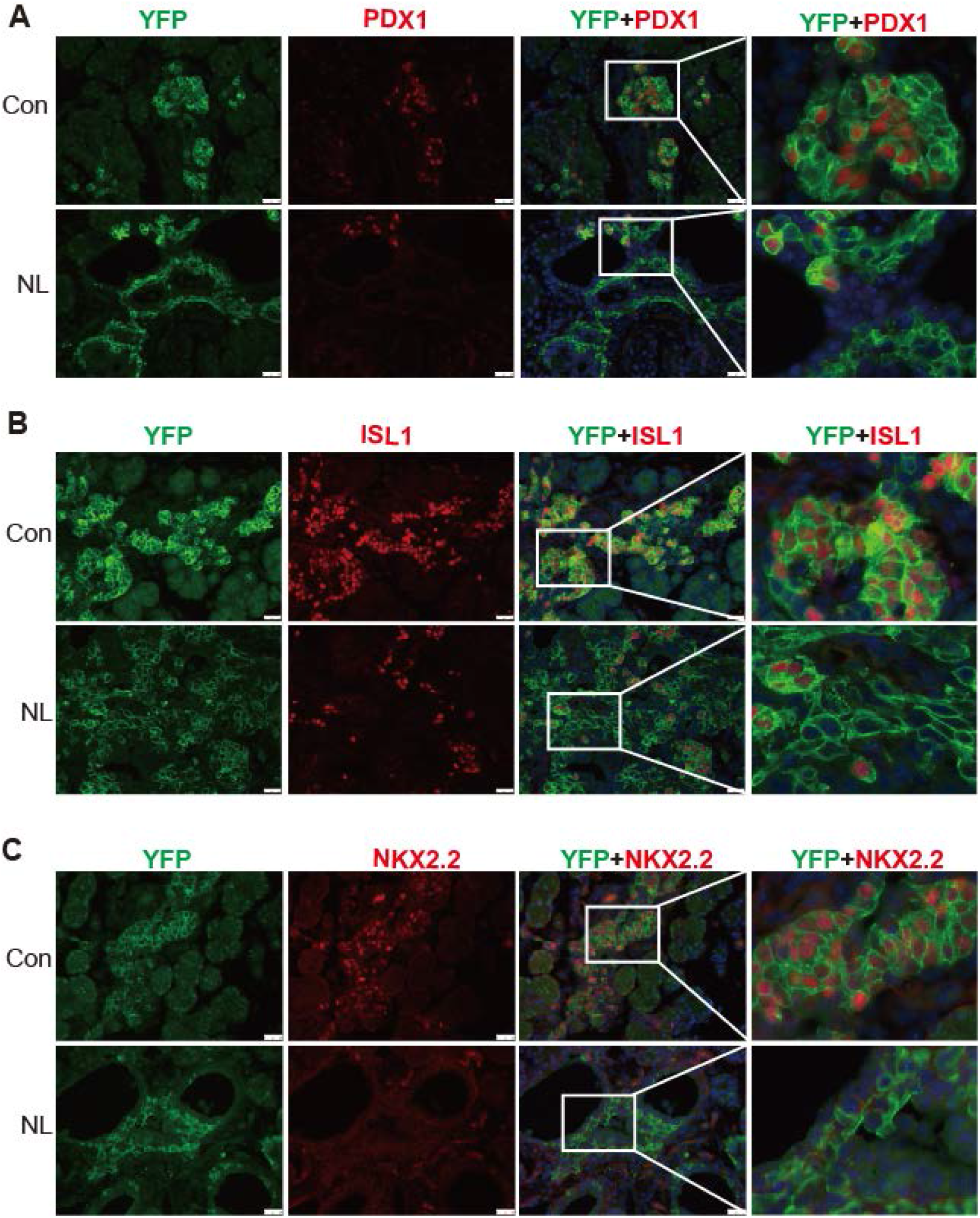
The differentiation of endocrine cells was blocked in NL pancreas at E16.5. A. Immunostaining of YFP and PDX1 in pancreatic tissue demonstrated that most of *Lats1/2* null YFP+ cells in NL mice were negative for PDX1 staining. B-C. Immunostaining of YFP and Islet 1 (ISL1, **B**) and NKX2.2 (**C**) showed that ISL1 and NKX2.2 were not expressed in the majority of YFP+ cells in NL pancreas.

### Loss of *Lats1&2* impairs epithelial-mesenchymal transition (EMT) in endocrine progenitor cells

During the endocrine formation process, several morphogenetic events coincide with cell differentiation including epithelial-to-mesenchymal transition (EMT) and delamination of differentiating endocrine cells during pancreatic development [15,16]. The new model proposed by Sharon et al. suggests that differentiating endocrine precursors cells undergo “leaving the cord” or “delamination” process with lower E-Cadherin (CDH1) expression[27]. How NGN3 mediates CDH1 downregulation and facilitates “leaving the cord” is unclear. We observed that YFP+ cells were connected to epithelial cords, but expressed lower CDH1 than the cord epithelial cells in control E16.5 pancreases (Figure 3A). YFP+ cells in the NL pancreas were connected to the epithelial cords and formed buds or sheaths similar to YFP+ cells in the control. However, they express CDH1 at a level as high as of that in the epithelial cord (Figure 3A). With respect to the few escapees as previously described in PDX1 expression, the NL pancreas, too, contained a few escapees marked by lower CDH1 expression. Interestingly, we found that the YFP+ cells in NL pancreases did not stay as “budding” or “sheath” as development continued, and returned to single epithelial cells at P1 (Figure S3). These data suggest that an active Hippo pathway is required for downregulation of CDH1 in *Ngn3* expressing progenitor cells, but is dispensable for “leaving cord” or “delamination”.

**Figure 3.**
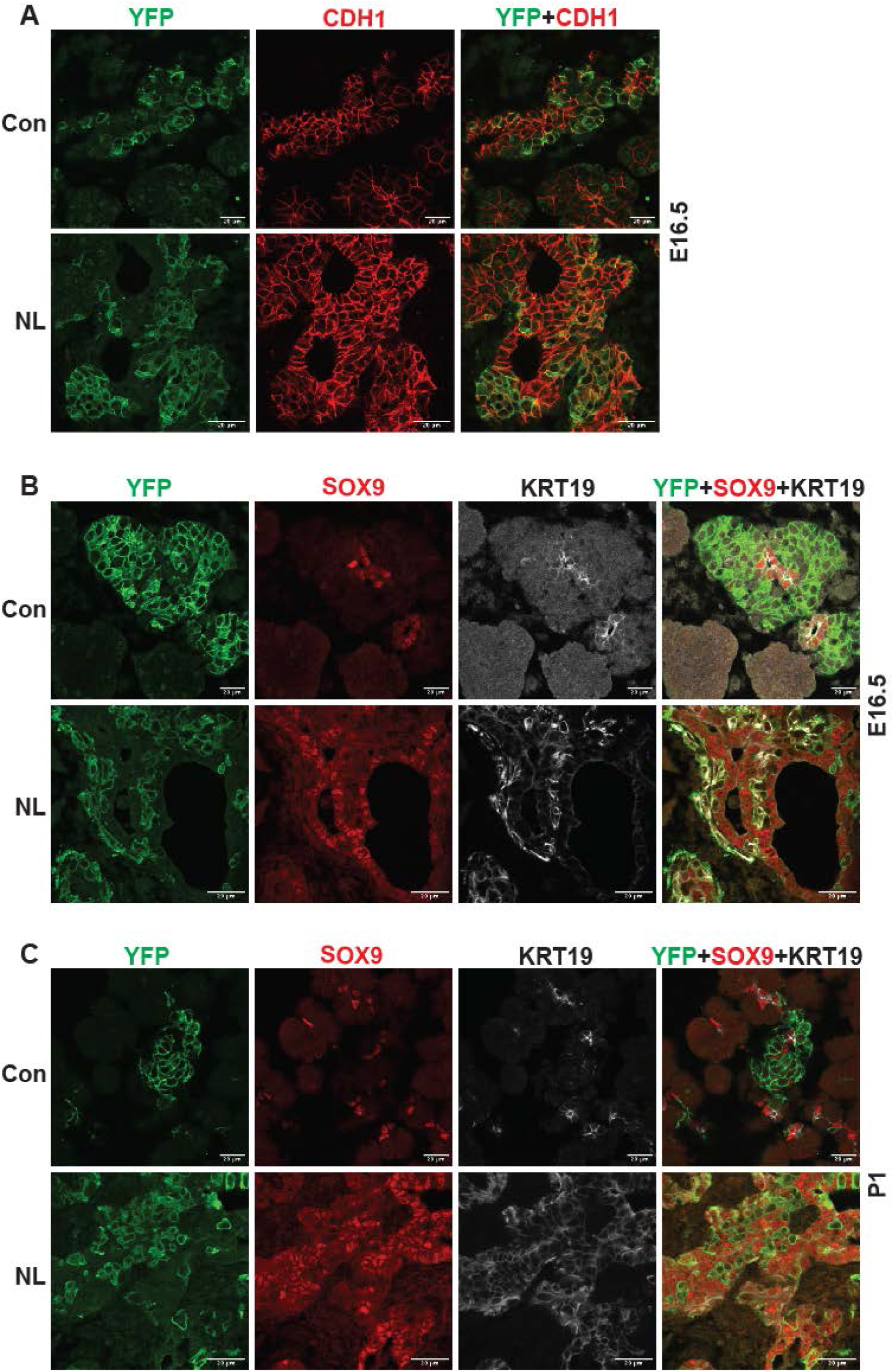
Loss of *Lats1&2* altered endocrine progenitor cell characteristics. A. CDH1 expression in YFP+ cells was low in control pancreas and considerably higher in NL pancreas at E16.5. B. In both control and NL pancreas, the YFP+ cells were negative for SOX9 staining at E16.5 and P1. C. KRT19 staining was negative in YFP positive cells in control pancreas but appeared to be positive in YFP+ cells of NL pancreas at both E16.5 and P1. Scale bar: 20 μm.

We have observed that the differentiation of *Lats1&2* null endocrine progenitor cells was blocked by lack of expression of ISL1 and NKX2.2. To expand on this, we interrogated the question of whether these progenitor cells returned to the ductal cell fate. To do so, we performed immunofluorescent staining and found that, similarly to the control pancreas, there was no overlap of YFP and SOX9 in the NL pancreas at E16.5 or P1 (Figure 3B). In addition, we examined another known ductal cell marker, cytokeratin 19 (KRT19) in the NL pancreas. We observed that KRT19 did not express in YFP+ cells in control pancreases, but was highly expressed in YFP+ cells of NL pancreases at E16.5 and P1 (Figure 3C). In fact, at E16.5, the expression of KRT19 in YFP+ cells of NL pancreases was even higher than the ductal cells, which later became more similar to ductal cells at the P1 stage (Figure 3C). These data suggest that an early effect of *Lats1&2* deletion in Ngn3+ cells is to turn on KRT19 expression, but not SOX9 expression, further indicating that KRT19 expression is not controlled by SOX9, but instead by YAP1.

### *Lats1&2* null cells recruit macrophages and induce pancreatitis-like phenotype in the developing pancreas

Following gene knockouts of *Lats1&2*, we observed that NL mice died before three weeks of age with a much smaller pancreas. Histological analysis showed acinar atrophy and disorganized ductal expansion in the NL pancreas, which are frequently associated with pancreatitis [28]. Our previous study involving acinar-specific deletion of *Lats1&2* (using *Ptf1a*^*CreER*^ model) has found that loss of *Lats1&2* first activates pancreatic stellate cells (PSC) [20]. We wondered whether a similar mechanism happened in the NL pancreas. Immunostaining showed that there was a significant increase of Vimentin-positive mesenchymal cells in the NL pancreas at E16.5 (Figure 4A). Those expanded mesenchymal cells were closely associated with YFP+ *Lats1&2* null cells (Figure S4A). These mesenchymal cells were positive for activated PSC marker alpha smooth muscle actin (ACTA2) (Figure S4B). Next, we determined whether there was immune cell infiltration in the NL pancreas. We observed CD45+ immune cells with large cell bodies, similar to the size of epithelial cells, in the E16.5 pancreas. We then stained tissue with macrophage marker F4/80 and found that these CD45+ cells were macrophages. The number of macrophage was increased in NL mice at E16.5, especially in the ductal area (Figure 4B and 4D), and further dramatically increased at P1 in the whole pancreas (Figure 4C and 4D). Overall, NL pancreases contained significantly greater macrophage cell density at E16.5 and P1 in comparison to control pancreases in both the acinar and ductal regions of the pancreas (Figure 4D). Together, these results indicate that loss of *Lats1&2* in NGN3+ endocrine progenitor cells activates PSCs in the developing pancreas, and recruits immune cells continuously.

**Figure 4.**
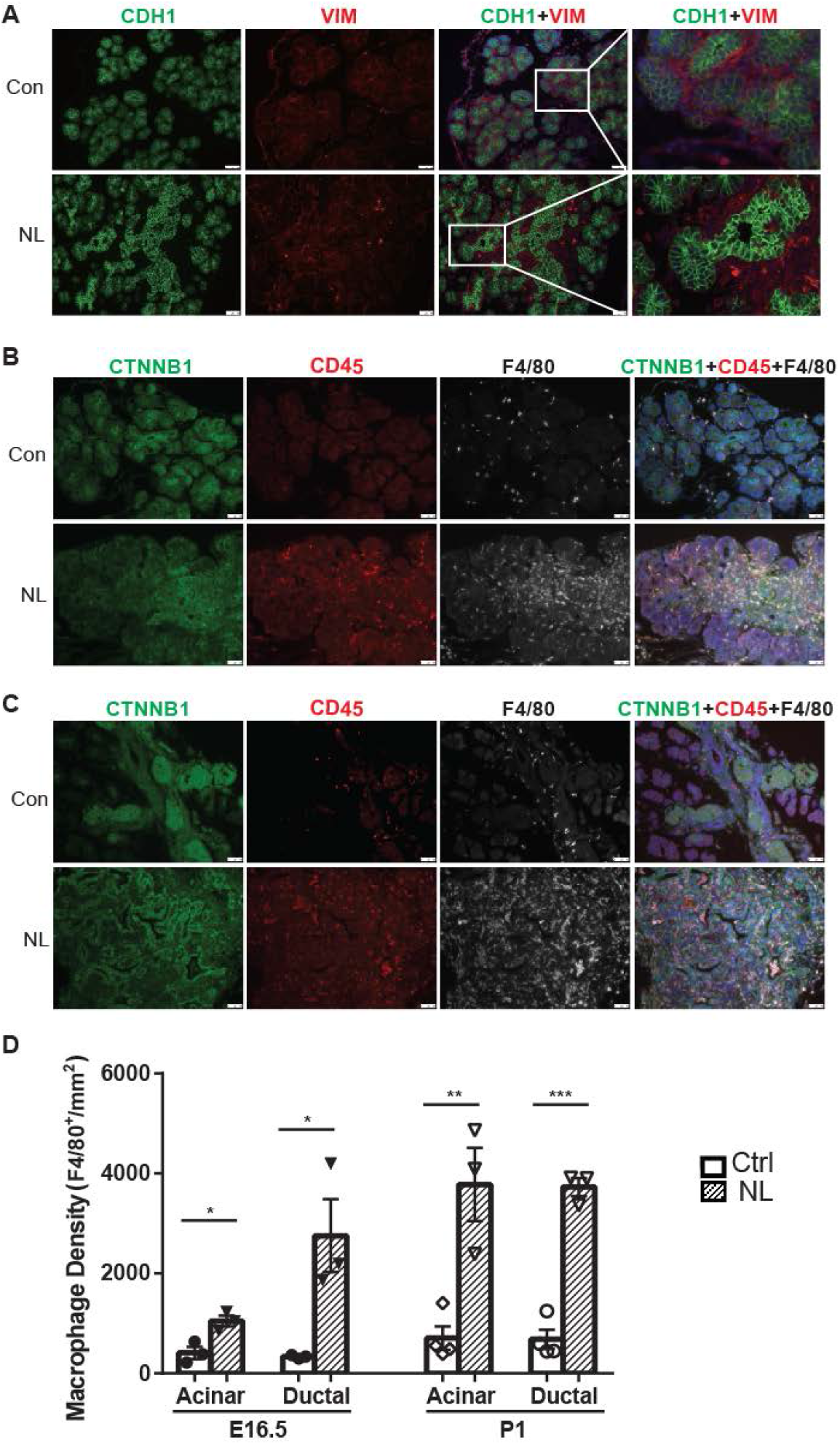
Loss of *Lats1&2* in endocrine progenitor cells was associated with an increased number of mesenchymal cells and macrophages. A. Vimentin positive staining was increased in NL pancreas at E16.5. Scale bar: 25μm. B-C. Immunostaining showed that the number of CD45 positive immune cells and F4/80 positive macrophage were significantly increased in NL pancreas at E16.5 (**B**), and became even more apparent at P1 (**C**). D. Quantification of F4/80 positive macrophage showed a significant increase in NL pancreas at both E16.5 and P1 (n=4). * p<0.05; ** p<0.01; *** p<0.001. Scale bar: 50μm.

### The expressions of YAP1/TAZ and their targets are elevated in *Lats1&2* null cells

Loss of *Lats1&2* in *Ngn3* expressing endocrine progenitor cells blocks the endocrine lineage from further differentiation. We hypothesized that uncontrolled downstream Hippo effectors, YAP1 and TAZ, mediate this phenotype. It was observed that YFP+ cells had no YAP1 expression in control pancreases at E16.5 (Figure 5A), suggesting YAP1 expression predominated in the epithelial cord region, which was consistent with the previous studies [9,17]. However, YAP1 expression was much higher in NL pancreases, with many YFP+ cells co-localizing with positive YAP1 nuclei staining despite a few escaped YFP+ cells lacking YAP1 expression (Figure 5A). We observed similar YAP1 staining pattern in P1 pancreases of NL mice (Figure S5A). We further co-stained NGN3 and YAP1 on E16.5 pancreases and found that, in the control pancreas, YAP1 was expressed in NGN3+ cells located in the epithelial cord, but was sequestered into the cytoplasm (Figure 5B). No YAP1 expression was detected in the cytoplasm or the nucleus in the NGN3+ cells after they left the epithelial cord. In comparison, much fewer NGN3+ cells, but more YAP1 positive cells, were observed in the NL pancreas, overall. In addition, nuclear YAP1 was co-stained with NGN3 in nuclei of cells residing in the epithelial cord of NL pancreases. Using qPCR, we then further quantified mRNA expression of YAP1 targets that we previously identified [20]. We found that the expressions levels of *Ankrd1, Cyr61*, and *TGFb1* had no significant changes, but *Ctgf, SPP1, Cxcl10*, and *Cxcl16* were significantly higher in the NL pancreas at P1 (Figure 5C). Immunofluorescent staining showed that SPP1 was expressed in ductal cells in Controlcontrol pancreas but was also positive in YFP+ cells in NL pancreas at E16.5 and P1 (Figure S5B). Together, these data demonstrate the increased expression of YAP1 and its target genes in *Lats1&2* null cells.

**Figure 5.**
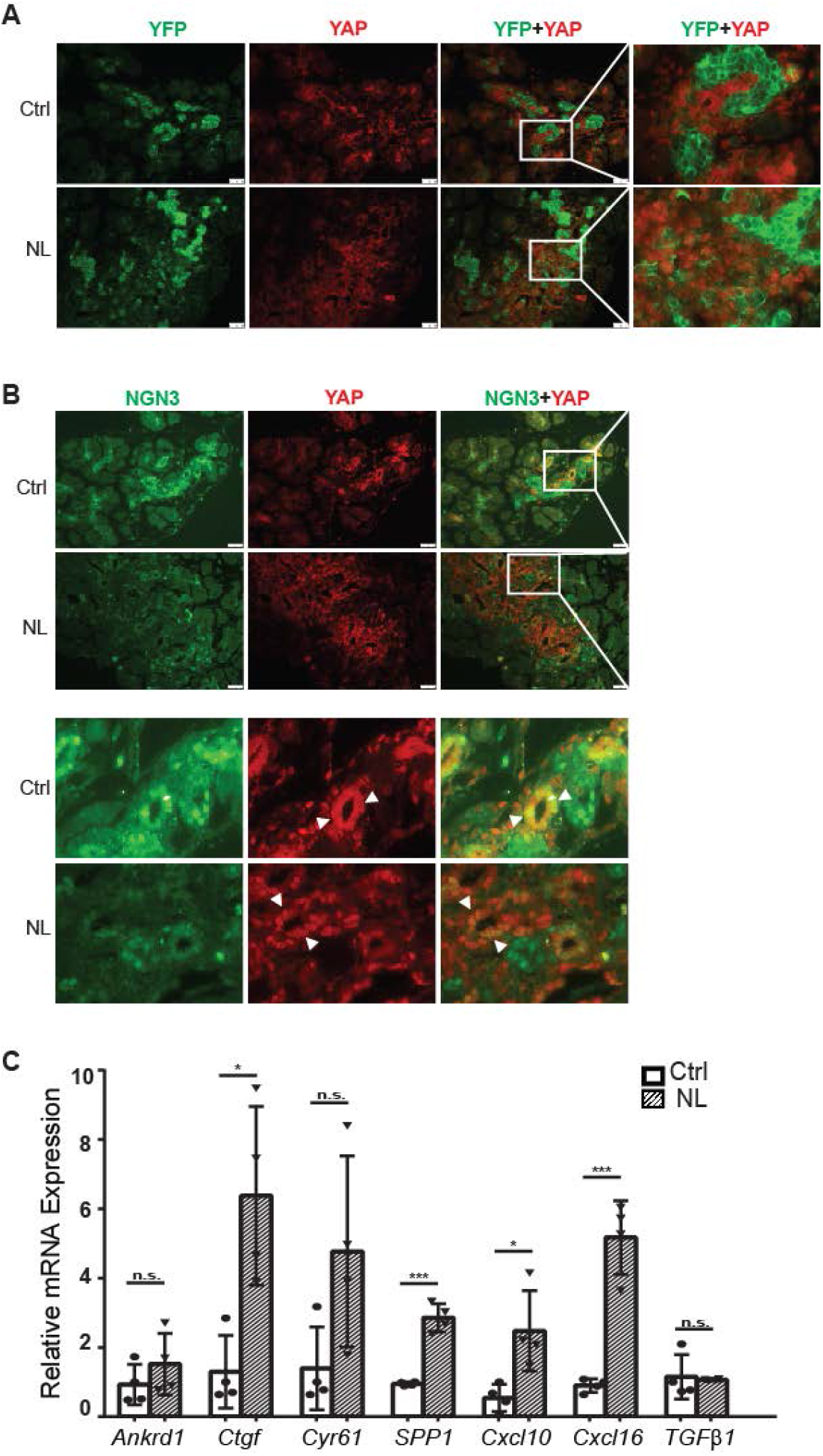
The expressions of YAP1/TAZ and their targets were increased in *Lats1&2* null cells. A. YAP1 was not expressed in YFP+ endocrine cells in control pancreas, but was found to be highly expressed in NL pancreas at E16.5. B. YAP1 was sequestered to the cytoplasm in newly born NGN3 positive cells (arrow head) located in the epithelial cord in control pancreas, while co-staining of YAP1 with NGN3 was observed in nuclei in epithelial cord of NL pancreas. C. The mRNA levels of YAP1 targets *Ctgf, SPP1, Cxcl10*, and *Cxcl16* were significantly increased in NL pancreas at P1 (n=4). * p<0.05; ** p<0.01; *** p<0.001. Scale bar: 50μm.

### YAP1/TAZ are downstream effectors of *Lats1&2* in regulating endocrine specification and differentiation

To further demonstrate that YAP1 and TAZ are the downstream effectors of *Lats1&2* in endocrine lineage development, we performed gene knockouts of *Yap1* and *Taz* (*Wwtr1*) in addition to *Lats1&2* in NGN3+ endocrine progenitor cells. To this end, we intercrossed *Ngn3*^*Cre*^*Lats1*^*fl/fl*^*Lats2*^*fl/+*^ with *Yap1*^*fl/fl*^*Taz*^*fl/fl*^ mice for several generations to obtain the following gene knockout mice models: *Ngn3*^*Cre*^*Lats1*^*fl/fl*^*Lats2*^*fl/+*^*Yap1*^*fl/fl*^*Taz*^*fl/fl*^ (named as NTY where *Lats2* is heterozygous and *Yap1/Taz* are null), *Ngn3*^*Cre*^*Lats1*^*fl/fl*^*Lats2*^*fl/fl*^*Yap1*^*fl/+*^*Taz*^*fl/fl*^ (named as NLT where *Yap1* is heterozygous but *Taz* is null), and *Ngn3*^*Cre*^*Lats1*^*fl/fl*^*Lats2*^*fl/fl*^*Yap1*^*fl/fl*^*Taz*^*fl/fl*^ (named as NLTY with quadruple null *Lats1&2, Yap1, and Taz*). First, we found that NTY mice were normal with no defect in the pancreas, suggesting that YAP1/TAZ are no longer needed in *Ngn3* expressing cells for endocrine lineage development. Because NL mice showed an obvious defect in the postnatal P1 pancreas, we focused on and analyzed the P1 pancreases of NLT and NLTY mice. Histological analysis showed that, unlike NL pancreases, NLTY pancreases were similar to control pancreases. However, NLT pancreases still had some abnormality in the somewhat abnormal ductal region, suggesting incomplete rescue when YAP1 remained expressed (Figure S6A). We further analyzed endocrine cells by staining YFP, INS, and Somatostatin (SST), and observed that only a small fraction of YFP+ cells, scattered throughout the NL pancreas, were INS and SST positive (Figure 6A and S6A). However, similar to control pancreases, most YFP+ cells in NLT and NLTY pancreases were INS and SST positive, suggesting that reduced levels of YAP1/TAZ rescued the *Lats1&2* null phenotype. However, we still observed some ductal-like YFP+ cells in NLT pancreases, again suggesting incomplete rescue compared to NLTY pancreases (Figure 6A). Furthermore, we analyzed immune cell infiltration and found that there were much fewer macrophages in both NLT and NLTY pancreases compared with NL pancreases (Figures 6B and 4C). In addition, quantification analysis showed that macrophage density is slightly elevated in NLT and NLTY pancreases compared to control pancreases, but not statistically significantly (Figure 6C). Taken together, our results demonstrate that YAP1/TAZ are the downstream effectors of *Lats1&2* during endocrine differentiation and a tight control of their activities is required for normal pancreatic development.

**Figure 6.**
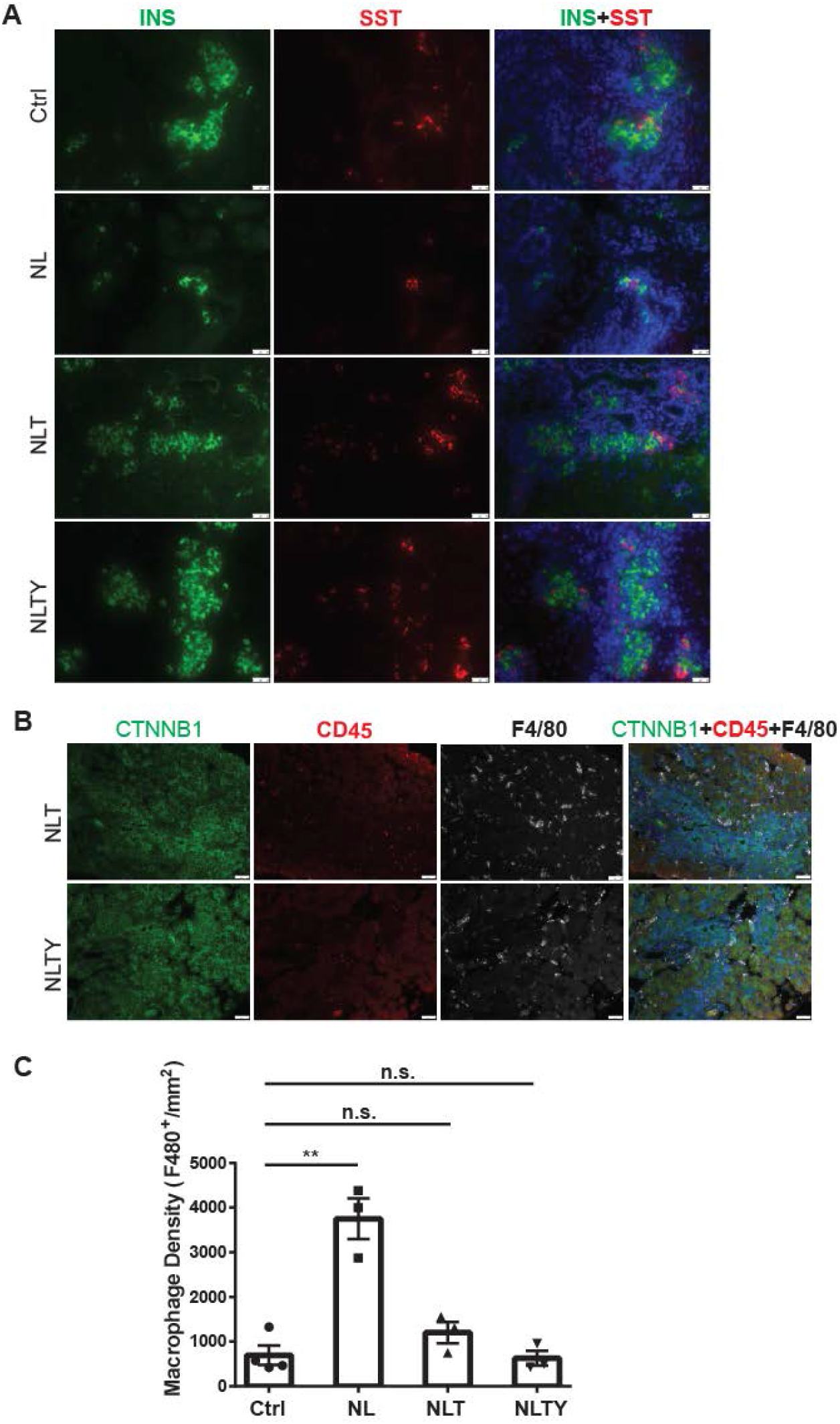
Removal of YAP1/TAZ rescued the defect in endocrine specification and differentiation in NL pancreas. A. Co-staining of Insulin (INS) and Somatostatin (SST) showed that onlysmall cell clusters were positive for INS or SST in NL pancreas, whereas the islets in NLT and NLTY pancreata were INS or SST positive, more closely resembling the control islets. Scale bar: 25μm. B. The numbers of immune cells were much less in both NLT and NLTY pancreata compared to NL pancreas. C. Quantification analysis showed that while the macrophage density of NL pancreas was significantly higher compared to control pancreas, there was no significant difference in macrophage density between both NLT and NLTY pancreas and control pancreas (n=4). n.s. p>0.05; ** p<0.01. Scale bar: 50μm.

### *Lats1&2* are dispensable for pancreatic β-cell proliferation and function

The established role of Hippo in regulating cell proliferation prompted us to investigate whether inactivation of *Lats1&2* affects pancreatic β-cell proliferation and function. To address this, we deleted *Lats1&2* in pancreatic β-cells using the mouse insulin promoter 1 driven CreER (*MIP1*^*CreER*^, named as ML). We first tested whether *Lats1&2* are necessary for pancreatic β-cell function at the adult stage (Figure 7A). The mice were subjected to five-day Tamoxifen (TAM) injection at 180 mg/kg/day, and pancreases were collected four weeks after TAM injection for histological analysis. Normal pancreatic architecture and similar endocrine cell mass were observed between control and ML pancreases through H&E staining (Figure 7B). To confirm the β-cell specific *Lats1&2* deletion efficiency, we performed western blot analysis and were unable to detect LATS1&2 protein in islets of ML mice (Figure 7C). To further test the influence of *Lats1&2* deficiency on β-cell function, we performed glucose tolerance tests (GTT). Results indicate that ML mice displayed a normal GTT profile when compared to control mice (Figure 7D).

**Figure 7.**
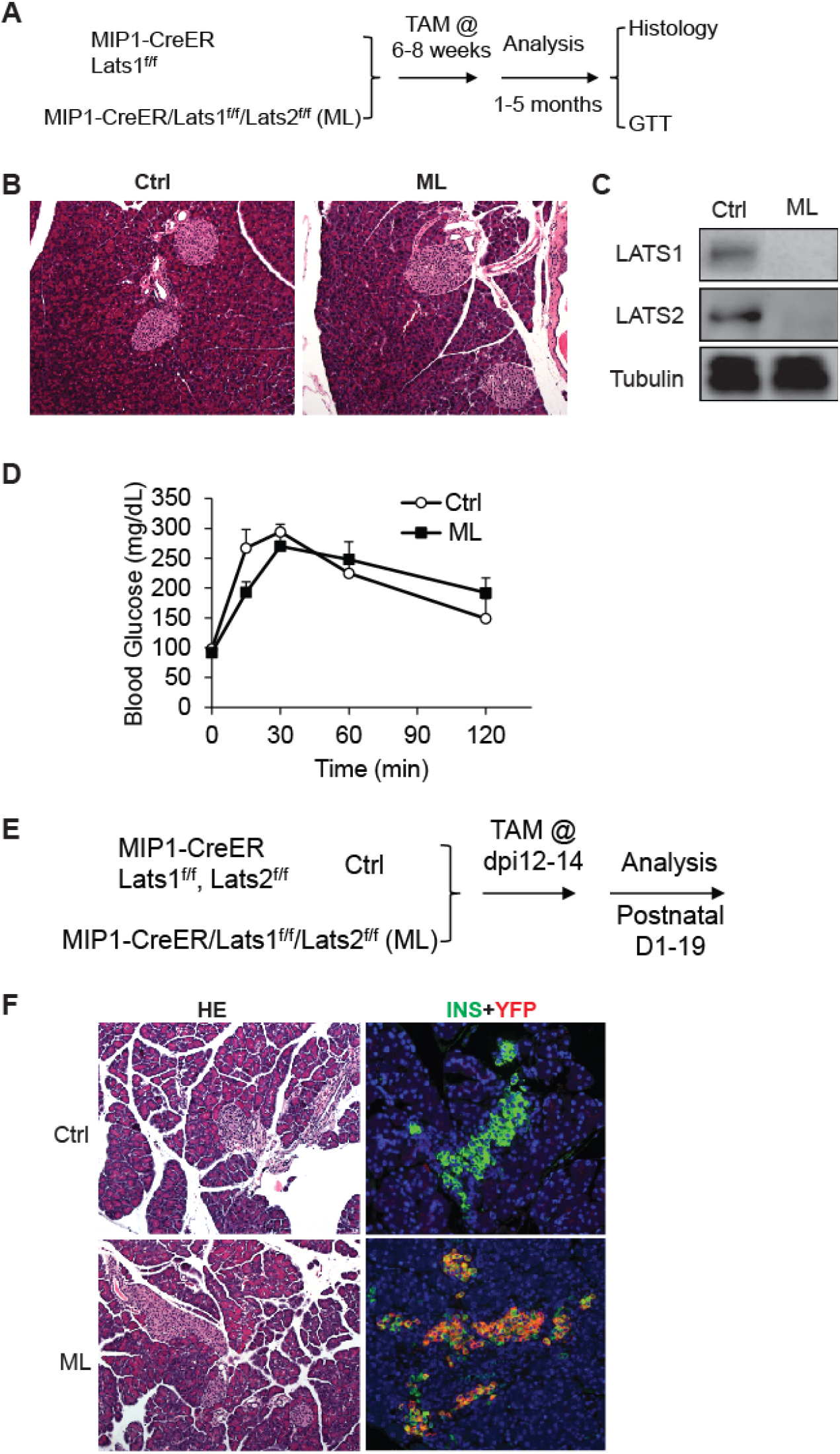
*Lats1&2* are dispensable for pancreas β-cell proliferation and function. A. Schematic strategy of deleting *Lats1&2* in adult β-cells using *MIP1*^*CreER*^ to generate ML mice. B. H&E staining showed normal pancreatic architecture and similar pancreatic islet size between control and ML pancreata. C. LATS1&2 protein levels were reduced in pancreatic islets of ML mice. D. The ML mice showed a normal GTT compared with control mice. (Ctrl, n=6, ML, n=5). E. Schematic strategy of deleting *Lats1&2* in embryonic β-cells. F. H&E staining showed normal pancreas structure between control and ML mice. Most YFP+ cells expressed INS in P1 ML pancreas.

Next, we tested whether *Lats1&2* is required for embryonic β-cells. To this end, we injected the pregnant mice carrying *MIP1*^*CreER*^*Lats1&2*^*fl/fl*^*Rosa26*^*YFP*^ offspring with tamoxifen at E12 (Figure 7E). The tamoxifen treatment induced the majority Insulin+ β-cells to express YFP at P1, showing an efficient gene deletion by this procedure (Figure 7F). However, the *Lats1&2* deletion did not affect pancreatic architecture and endocrine pancreas morphology (Figure 7G). Together, these findings suggest that *Lats1&2* are dispensable for pancreas β-cell proliferation and function.

## Discussion

Recent studies have revealed that the Hippo signaling pathway and its effectors are essential for pancreatic development and function. Multiple Hippo pathway genes have been investigated through specific deletion at embryonic and adult stages using different pancreatic-specific Cre lines in genetically engineered mice models [9–11,20]. With the early embryonic deletion of *Lats1&2* using *Pdx1*-early Cre, the developing pancreas loses cell polarity and subsequent undergoes failure of epithelial expansion/branching, indicating that early pancreatic morphogenesis requires a properly functioning Hippo signaling pathway. However, the exact role of the Hippo signaling pathway in endocrine lineage specification has been unclear. Using mice with *Ngn3* driven Cre, we have revealed that the tight regulation of the Hippo pathway is required for endocrine specification and differentiation. Loss of *Lats1&2* in endocrine progenitor cells blocks their further differentiation, resulting in much smaller pancreatic islets and fewer hormone-producing cells. Further deletion of YAP1/TAZ rescues the observed endocrine defects in the *Lats1&2* null pancreas, suggesting that YAP1/TAZ transcriptional activities must be tightly controlled by LATS1&2 for endocrine lineage development. Our findings, therefore, expand our understanding of the physiological functions of the Hippo pathway (Figure 8).

**Figure 8.**
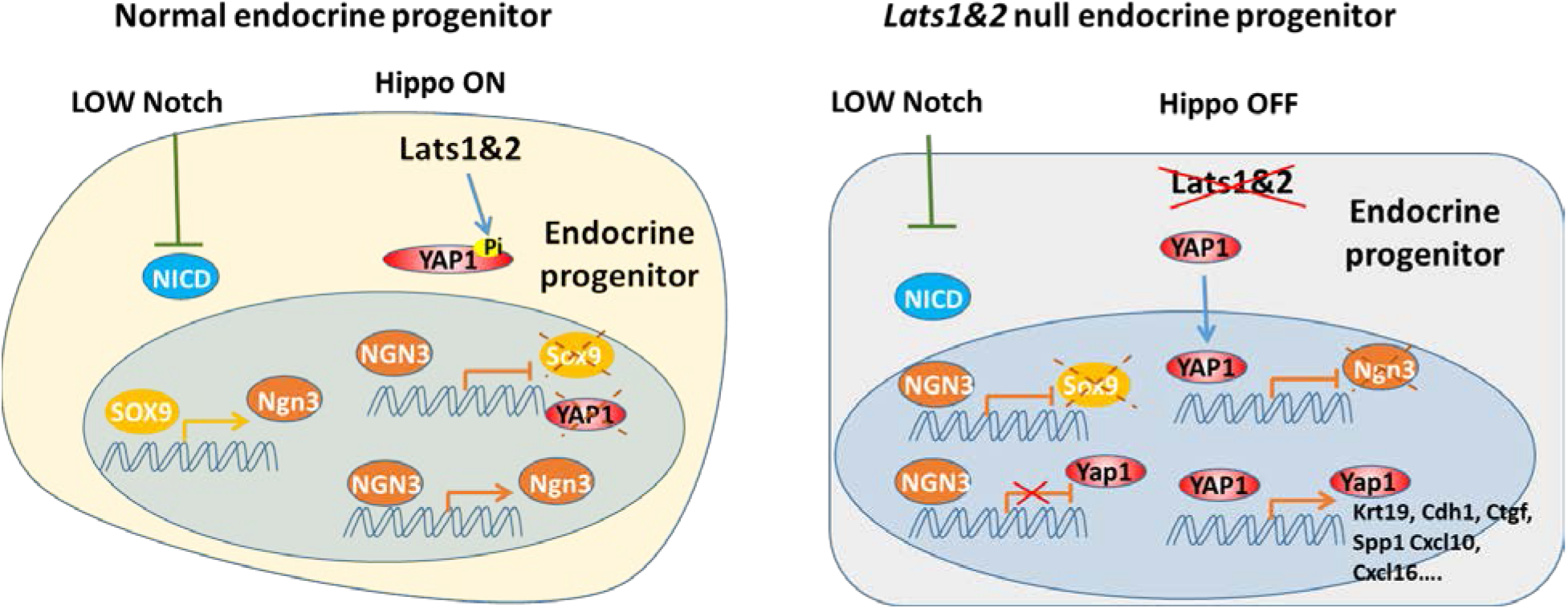
During endocrine progenitor specification, the Hippo pathway is required to sequester YAP1 in the cytosol and allow NGN3 to positively regulate its own expression and suppress *Sox9* and *Yap1* expression. Loss of *Lats1&2* leads to YAP1 activation which suppresses *Ngn3* expression and induces the expression of YAP1 targets.

Accumulating evidence has shown that Hippo plays dominant roles in the developing exocrine pancreas, but not in the developing endocrine pancreas [9,10,17]. First, deleting *Mst1&2* with an inducible acinar cell-specific Cre or overexpressing YAP1 in the pancreas reproduced the *Mst1&2* whole pancreas epithelial knockout phenotype—acinar atrophy, ductal expansion, and pancreatitis-like phenotype [9]. In addition, YAP1/TAZ were undetectable in both embryonic endocrine cells and adult endocrine cells [10,17]. Overexpression of *Ngn3* silenced YAP1/TAZ at the transcriptional level [17]. Consistent with these findings, we also found that embryonic NGN3+ cells are YAP1 negative. However, newly formed NGN3+ cells still exhibit YAP1 expression, but mainly in the cytosol, suggesting that YAP1 is controlled at the post-translational level, most likely by LATS1&2, the canonical Hippo pathway. *Ngn3* is a master regulator of endocrine differentiation and will turn on downstream genes including *Islet1* and *Nkx2*.*2*. In our *Lats1&2* null cells, YAP1 remains high in *Ngn3* expressing cells. This blocks the endocrine progenitor cells from expression of downstream transcription factors, such as *Islet1* and *Nkx2*.*2*, suggesting that high YAP1 blocks endocrine differentiation at a very early stage. Our data showed that NGN3 alone is not sufficient to shut off YAP1 expression. It requires active LATS1&2 to facilitate YAP1/TAZ sequestration outside of cell nuclei. Deletion of *Lats1&2* in NGN3 expressing cells disrupts NGN3’s ability to suppress YAP1/TAZ expression. This suggests that YAP1 is upstream of NGN3 and may regulate its own expression. Using human embryonic pancreases and embryonic-stem-cell-derived progenitors, Cebola et al. have found that YAP1 coactivator TEAD1 is a vital component of the combination of transcription factors that activates both stage and lineage-specific pancreatic enhancers of multipotent progenitor cells [29]. When we checked the list of TEAD1 binding peaks from Cebola’s data set, we found that TEAD1 binds to the YAP1 promoter region. Further investigation of YAP1 autoregulation and how NGN3 shuts down YAP1 expression during endocrine differentiation is warranted.

During endocrine pancreas formation, *Ngn3* has to achieve high expression levels for endocrine commitment[14,30]. High *Ngn3* initiates a stepwise differentiation process including epithelial-to-mesenchymal transition (EMT) and delamination of differentiating endocrine cells. The new inclusive model proposed by Sharon et al., using single cell sequencing and immunostaining, suggests that islets form in a budding process [27]. Bipotent progenitor cells with a sufficiently high level of NGN3 may leave the epithelial cord, but still remain attached to the cord. Sharon et al. suggested that differentiating endocrine precursors do not undergo EMT, but still express E-cadherin (CDH1) throughout the differentiation process. However, they observed downregulation of CDH1 during endocrine lineage formation, as previously found [15]. “Leaving the cord” or “delamination” has been suggested to follow *Ngn3* expression. How NGN3 mediates CDH1 downregulation and facilitates this process is unclear. We had similar observations in that NGN3 expressing YFP+ cells were connected with epithelial cords but with lower CDH1 staining in control E16.5 pancreas. However, YFP+ cells in the NL pancreas were connected to the cords and formed buds or sheaths similar to the control YFP cells, but they expressed CDH1 as highly as in the epithelial cord. Our findings suggest that high YAP1 activity maintains high CDH1 expression irrespective of *Ngn3* expression. We also found that TEAD1 binds to the *Cdh1* promoter region in Cebola’s data set[29], indicating that YAP1 maintains CDH1 expression. However, it is unclear whether loss of YAP1 expression is sufficient to downregulate CDH1 in the endocrine lineage. Nevertheless, our findings suggest that an active Hippo pathway is required for the downregulation of CDH1 in *Ngn3* expressing progenitor cells. Notch and Hippo signaling pathways are tightly regulated to govern endocrine lineage determination [31,32]. High Notch signaling without Hippo generates high HES1 and YAP1, which cooperate to suppress *Ngn3* expression, leading to ductal cell fate. Low Notch signaling with Hippo activity results in *Ngn3* expression mediated by SOX9. The role of SOX9 in endocrine lineage differentiation is complex. SOX9 expresses in the Notch-responsive bipotent progenitors and committed ductal cells. It is suppressed by NGN3 in committed endocrine progenitors[33]. We found that newly born NGN3+ cells sequestered YAP1 out of nuclei by the Hippo pathway. When NGN3+ cells grow out of the epithelial cord, YAP1 expression is completely turned off. Loss of *Lats1&2* results in high YAP1 in the nuclei of newborn NGN3+ cells, which blocks NGN3’s ability to turn on endocrine lineage gene expression. Interestingly, *Sox9* has been downregulated in these cells, suggesting that a short duration of *Ngn3* expression has shut off *Sox9* expression and that YAP1 cannot reactivate *Sox9* expression. Mamidi et al. have shown that overexpression of YAP1 in progenitor cells can expand the ductal compartment [31]. In this setting, *Ngn3* has not been turned on and SOX9 is high, implying that ductal expansion is probably mediated by both SOX9 and YAP1. In our NL mice, *Ngn3* and Cre-recombinase expression are induced simultaneously. Subsequently, *Lats1&2* are deleted and result in YAP1 activation. In these mutant cells, NGN3 has already turned off SOX9 expression. However, the ductal marker KRT19 is as high as in the ductal cells, suggesting that KRT19 most likely is regulated by YAP1, not by SOX9. Interestingly, a TEAD1 binding peak was found in *Krt19* gene promotor region in Cebola’s data set[29]. Although the mutant cells have high CDH1 and KRT19, and low SOX9, they still leave the epithelial cord and form the sheath, suggesting that “leaving the cord” is mediated by NGN3. However, high YAP1 activity in the mutant cells does not support *Ngn3* expression, resulting in no continuous growth of the sheath. The bulged-out cells return to the epithelial cord. Interestingly, SOX9 remains low in these mutant cells even if they return to simple epithelial cells.

Using human embryonic stem cells, Rosado-Olivieri et al. have showed that overexpression of constitutively active YAP1 impairs endocrine differentiation while inhibition of YAP1 can generate improved insulin-secreting cells [34]. This is consistent with our *in vivo* findings. Interestingly, others have found that overexpression of an active form of YAP1 greatly induces β-cell proliferation in adult human islets [17,35]. Our genetic models indicate that in adult β-cells, YAP1 expression has been silenced at the transcription level. Loss of *Lats1&2* has no effect on β-cell function. Direct increase of YAP1 expression may be required for expansion of β-cells in the adult pancreas. Thus, manipulating YAP1 level to increase β-cell number depends on cell types and developmental stage.

By acinar cell specific deletion of *Lats1&2*, we showed that PSC activation precedes macrophage activation and infiltration, resulting in extensive fibrosis [20]. In our NL pancreas, some PSC activation is visible, but macrophage infiltration is very distinct even at E16.5 and becomes more conspicuous at P1. Braitsch et al.’s study in which they deleted *Lats1&2* in the very early stage of pancreas development also had similar observations [11], i.e. the CD45+ cells surrounded the E11.5 mutant pancreas and infiltrated the E14.5 mutant pancreas. We have found that the chemokines *Cxcl10* and *Cxcl16* were upregulated in NL pancreases. These chemokines have been shown to recruit macrophages. We also found that those chemokines are upregulated in our acinar cell specific *Lats1&2* knockout pancreas, suggesting that YAP1/TAZ activation may generate signals to directly recruit macrophages to the tissue. The tissue-resident macrophages have been suggested to seed the tissue during embryonic development from the blood island of yolk sac and fetal hematopoietic cells in the liver [36,37]. However, the exact nature of tissue-resident macrophages and the signaling pathways that promote their migration and residency are still illusive. The fact that the macrophage recruitment can happen as early as E11.5 in the Hippo-off state implies that Hippo signaling may play an important role in recruiting tissue resident macrophages during organ development. Further in-depth investigation is required to test this possibility.

In conclusion, our study suggests that proper Hippo activity is required for the *Ngn3* driven differentiation program, further expanding our fundamental understanding of Hippo pathway participation in pancreatic endocrine development.

## Acknowledgments

The authors acknowledge Dr. Andrew Leiter and Dr. Seung Kim for kindly providing the *Ngn3*^*Cre*^ mouse line, Dr. Randy Johnson for kindly providing the *Lats1*^*fl/fl*^ and *Lats2*^*fl/fl*^ mouse line, and Dr. Eric N. Olson for kindly providing the *Yap1*^*fl/fl*^ and *Taz*^*fl/fl*^ mouse line. The authors thank Dr. Guoqiang Gu (Vanderbilt University) and Dr. Yi Xu for their critical comments.

## Funding Statement

Pei Wang is a CPRIT scholar. This work is supported by the Cancer Prevention and Research Institute of Texas (P. Wang, R1219) and NIDDK (P. Wang, R01DK110361). The P.W. group was supported by Cancer Prevention and Research Institute of Texas, the William and Ella Owens Medical Research Foundation, and National Cancer Institute (R21 CA218968, R01 CA237159). Michael Nipper and Xue Yin were supported by a pre-doctoral fellowship through CPRIT Research Training Award RP 170345; Jun Liu was supported by a post-doctoral fellowship through CPRIT Research Training Award RP140105. The funders had no role in study design, data collection and analysis, decision to publish, or preparation of the manuscript.

## Author Contributions

Conceptualization: Pei Wang

Data curation: Yifan Wu, Kunhua Qin, Kevin Lopez, Jun Liu, Janice Deng, Michael Nipper, Xue Yin, Logan Ramjit, Pei Wang

Formal analysis: Pei Wang, Yifan Wu, Kunhua Qin, Kevin Lopez, Jun Liu, Michael Nipper, Zhengqing Ye

Funding acquisition: Pei Wang

Methodology: Yifan Wu, Kunhua Qin, Kevin Lopez, Jun Liu, Michael Nipper.

Project administration: Pei Wang

Supervision: Pei Wang

Writing – original draft: Yifan Wu, Kunhua Qin, Pei Wang

Writing – review & editing: Kevin Lopez, Michael Nipper, Jun Liu, Pei Wang

## Competing interests

The authors have declared that no competing interests exist.

## Abbreviations

ACTA2: α-smooth muscle actin
Amy: Amylase
BSA: bovine serum albumin
CDH1: E-Cadherin
Cpa1: carboxypeptidase A1
ChrA: Chromogranin A
CD45: cluster of differentiation antigen 45
KRT19: cytokeratin 19
CTGF: connective tissue growth factor
GCG: Glucagon
GTT: glucose tolerance tests
HE: hematoxylin–eosin
IF: immunofluorescent staining
Ins1: Insulin 1
Ins2: Insulin 2
i. p.: intraperitoneal
Isl1: transcription factors Islet 1
NKX2.2: NK2 homeobox 2
Ki67: antigen identified by monoclonal antibody Ki 67
LATS1: large tumor suppressor 1
LATS2: large tumor suppressor 2
MST1&2: Ste-20-like protein kinases 1 and 2
Ngn3: Neurogenin 3
Pdx1: pancreatic and duodenal homeobox 1
PBS: phosphate-buffered saline
PFA: paraformaldehyde
PSC: pancreatic stellate cell
qPCR: quantitative PCR
RT: reverse transcription
SPP1: secreted phosphoprotein 1
TAM: tamoxifen
TAZ: transcriptional coactivator with PDZ binding motif
TEAD: TEA domain family member
Yap1: yes-associated protein 1
YFP: yellow fluorescent protein

## Notes

### Competing Interest Statement

The authors have declared no competing interest.

