## Supplementary material for "Hippo pathway-mediated YAP1/TAZ inhibition is essential for proper pancreatic endocrine specification and differentiation": NL-supplement: Supplemental data-1.pdf

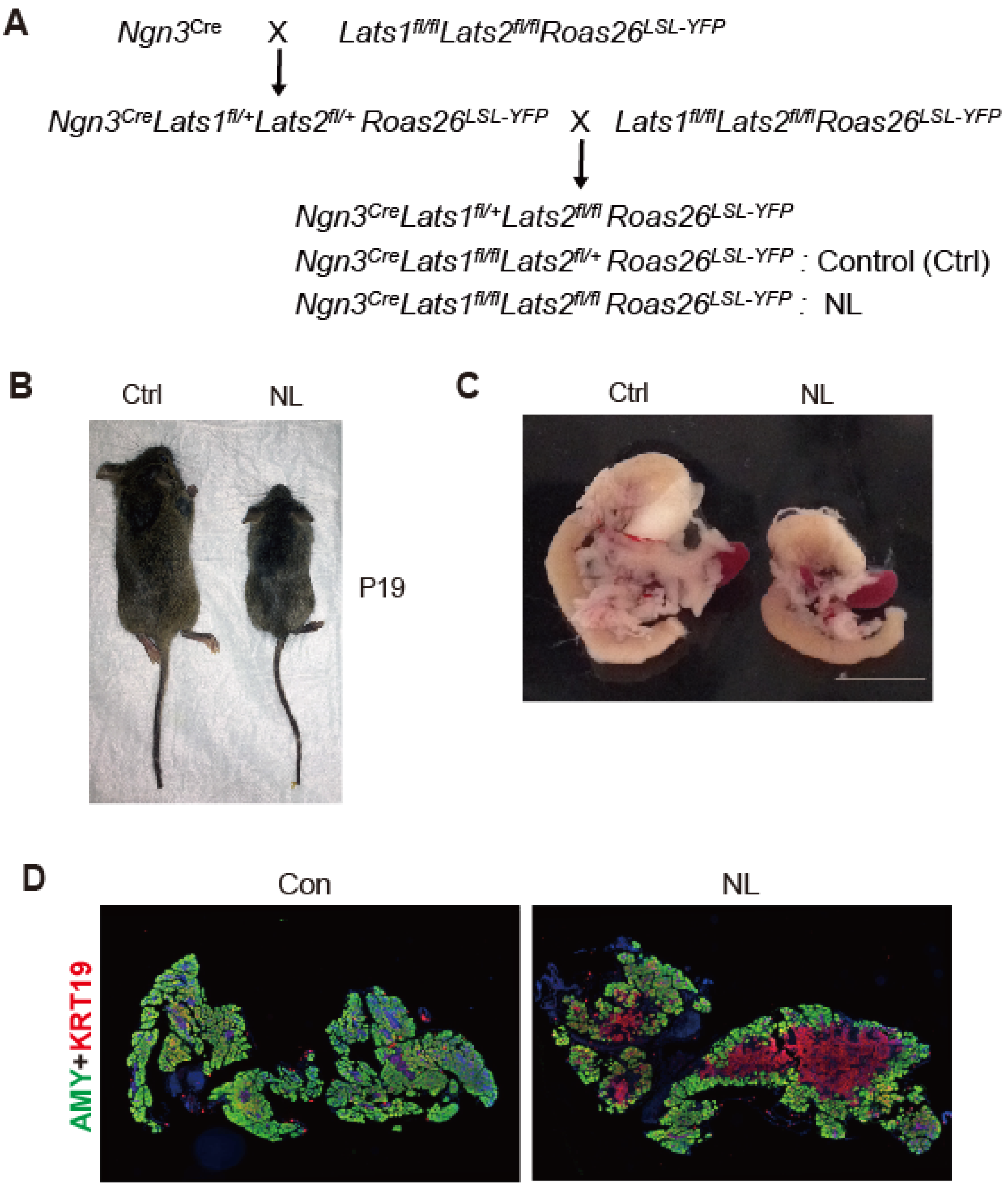


Supplemental Figure 1. Deletion of *Lats1/2* by *Ngn3*$^{Cre}$ perturbed pancreas differentiation. A. Schematic mating strategy of generating NL mice. B. NL mice were associated with smaller body sizes as compared to the littermate controls at P19. C. Macroscopic image of dissected P1 pancreas from control and NL mice. D. Architectural changes of NL pancreas at P1 were determined by immunostaining with Amylase (AMY) and KRT19.

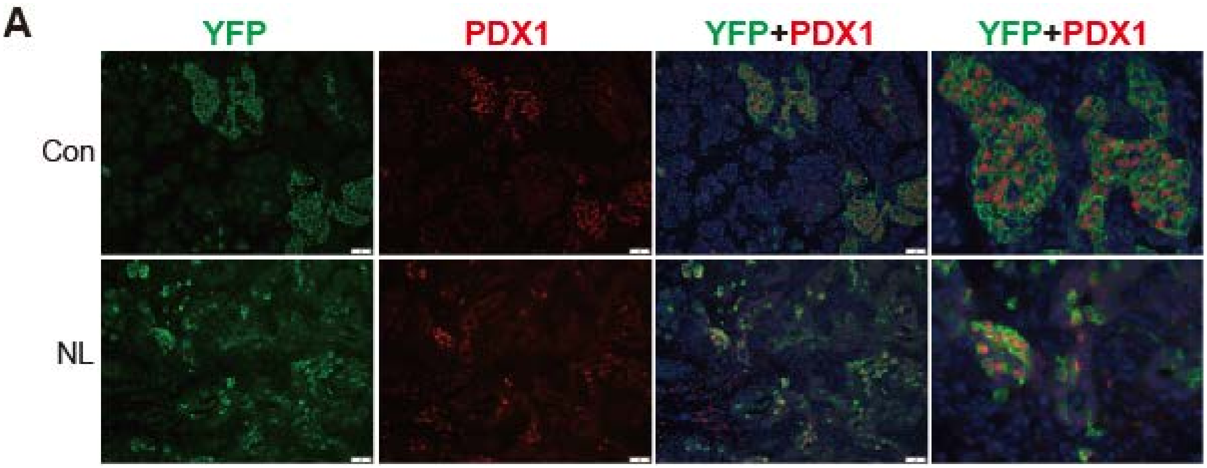


Supplemental Figure 2. The differentiation of endocrine cells was blocked in NL pancreas at P1. A. PDX1 was only expressed in a small fraction of YFP+ cells in the NL pancreas.

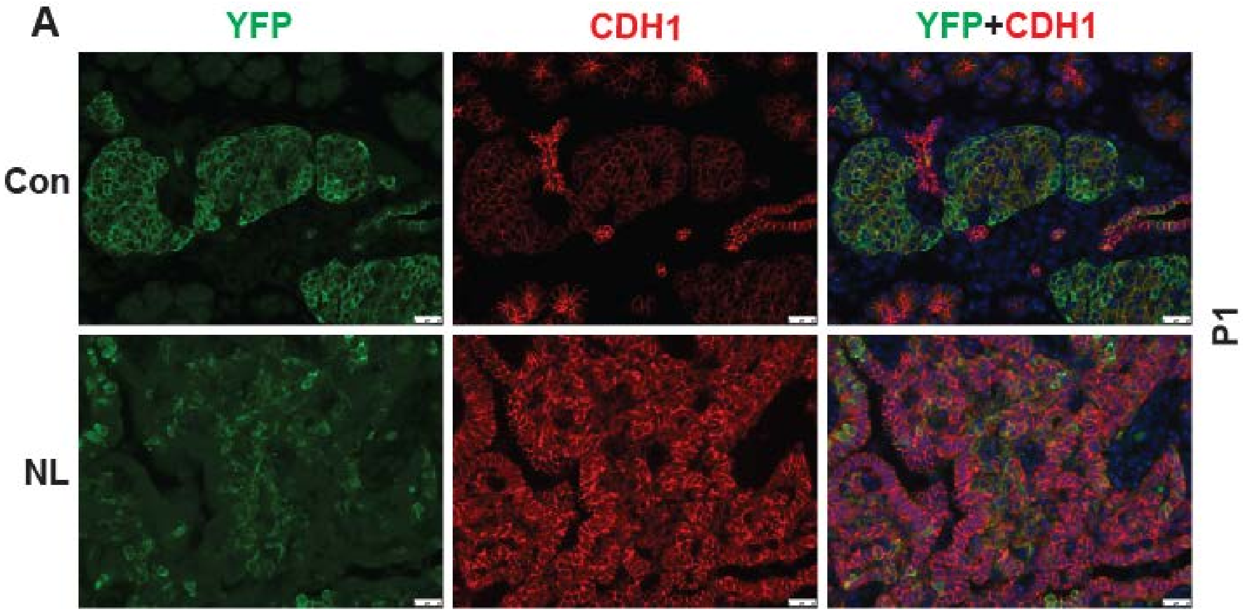


Figure S3. A. CDH1 expression in YFP+ cells was low in control pancreas and considerably higher in NL pancreas at P1. Scale bar: 25μm.

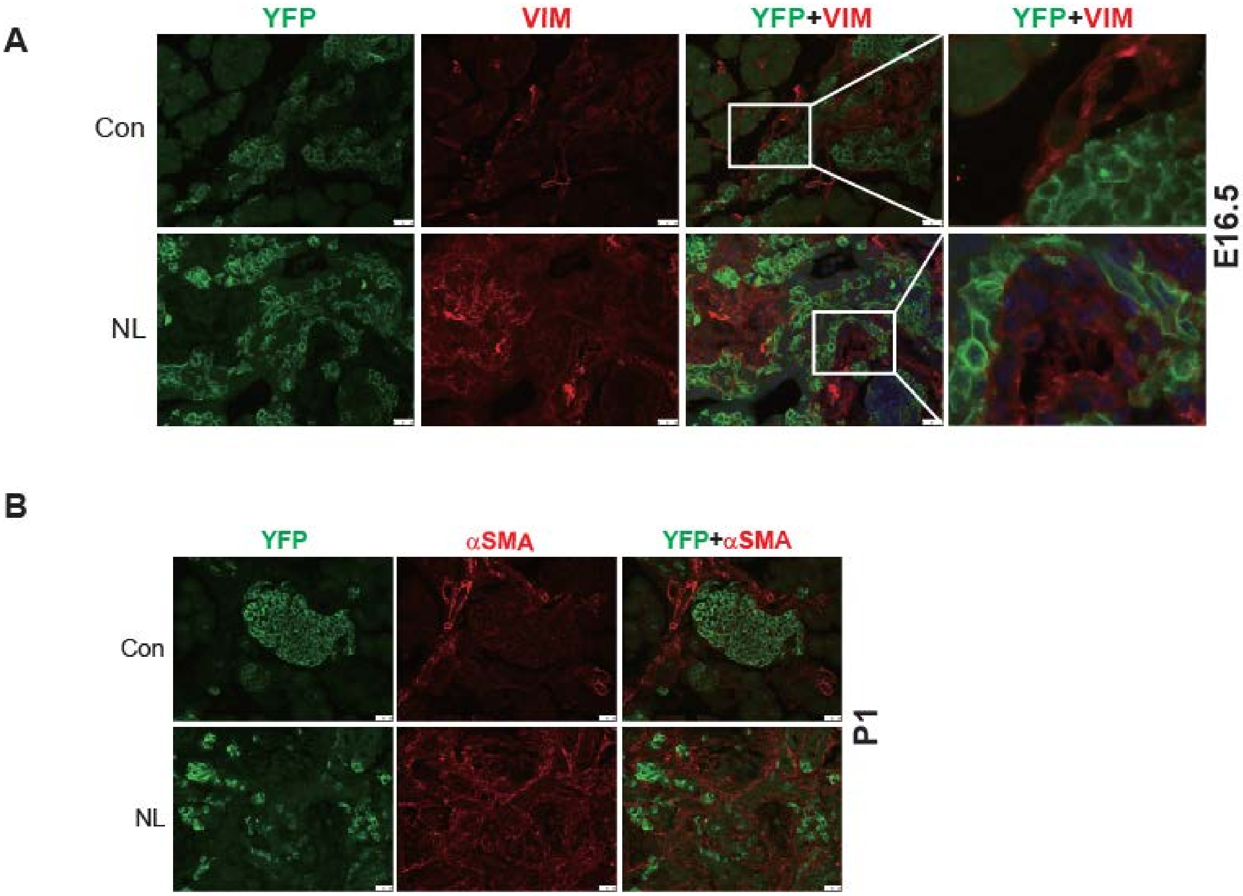


Figure S4. Loss of *Lats1&2* in endocrine progenitor cells was associated with increased mesenchymal cells. A. Immunostaining of YFP and Vimentin revealed the expansion of mesenchymal cells next to YFP+ *Lats1&2* null cells at E16.5. B. The mesenchymal cells were positive for αSMA in P1 NL pancreas. Scale bar: 25μm.

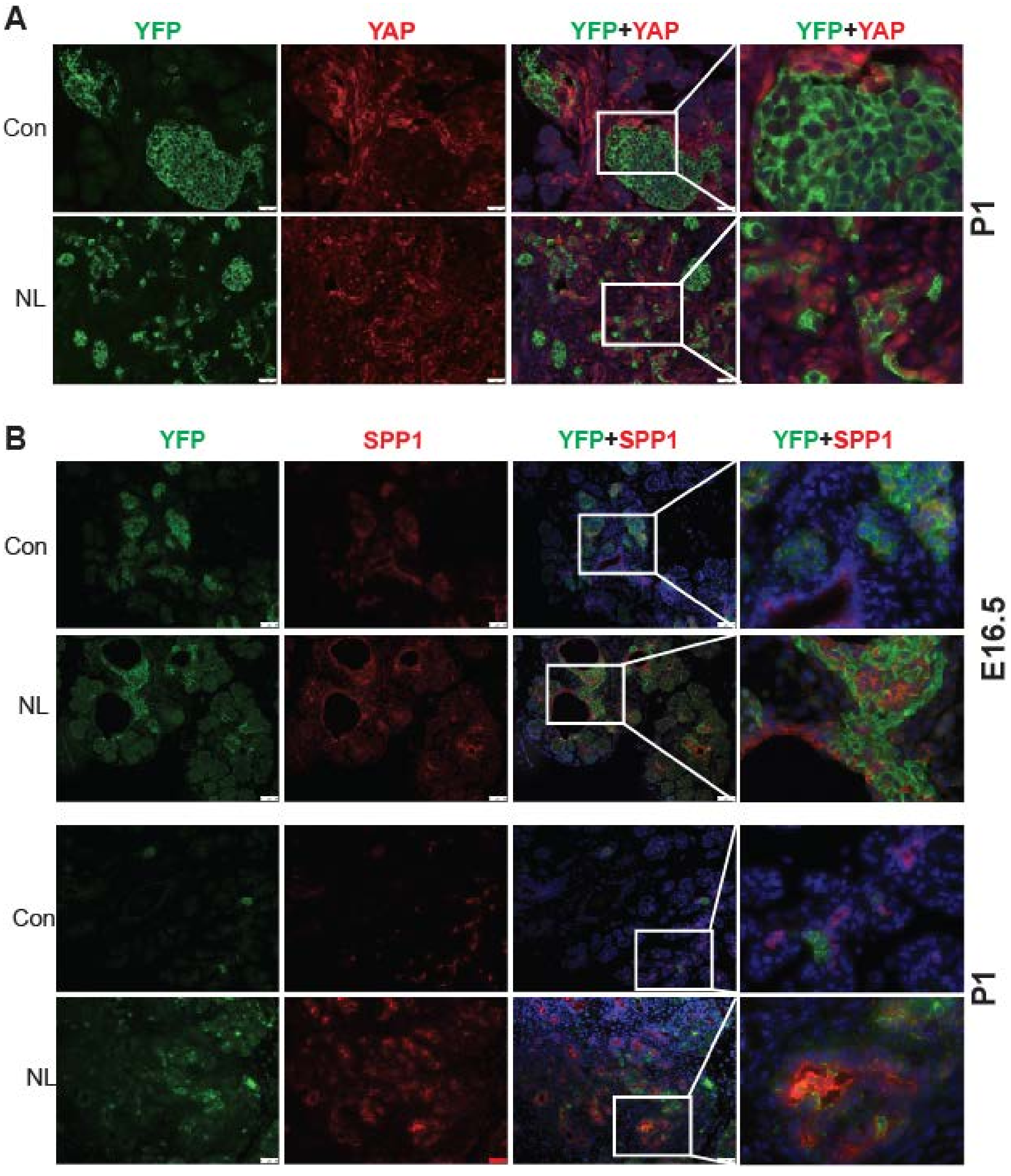


Figure S5. YAP1 and its targets were increased in *Lats1&2* null cells. A. YAP1 was not expressed in YFP+ endocrine cells in control pancreas, but was present in nuclei of YFP+ cells in NL pancreas at P1. B. SPP1 was expressed in YFP+ cells in NL pancreas, but not in control at E16.5 and P1. Scale bar: 50μm.

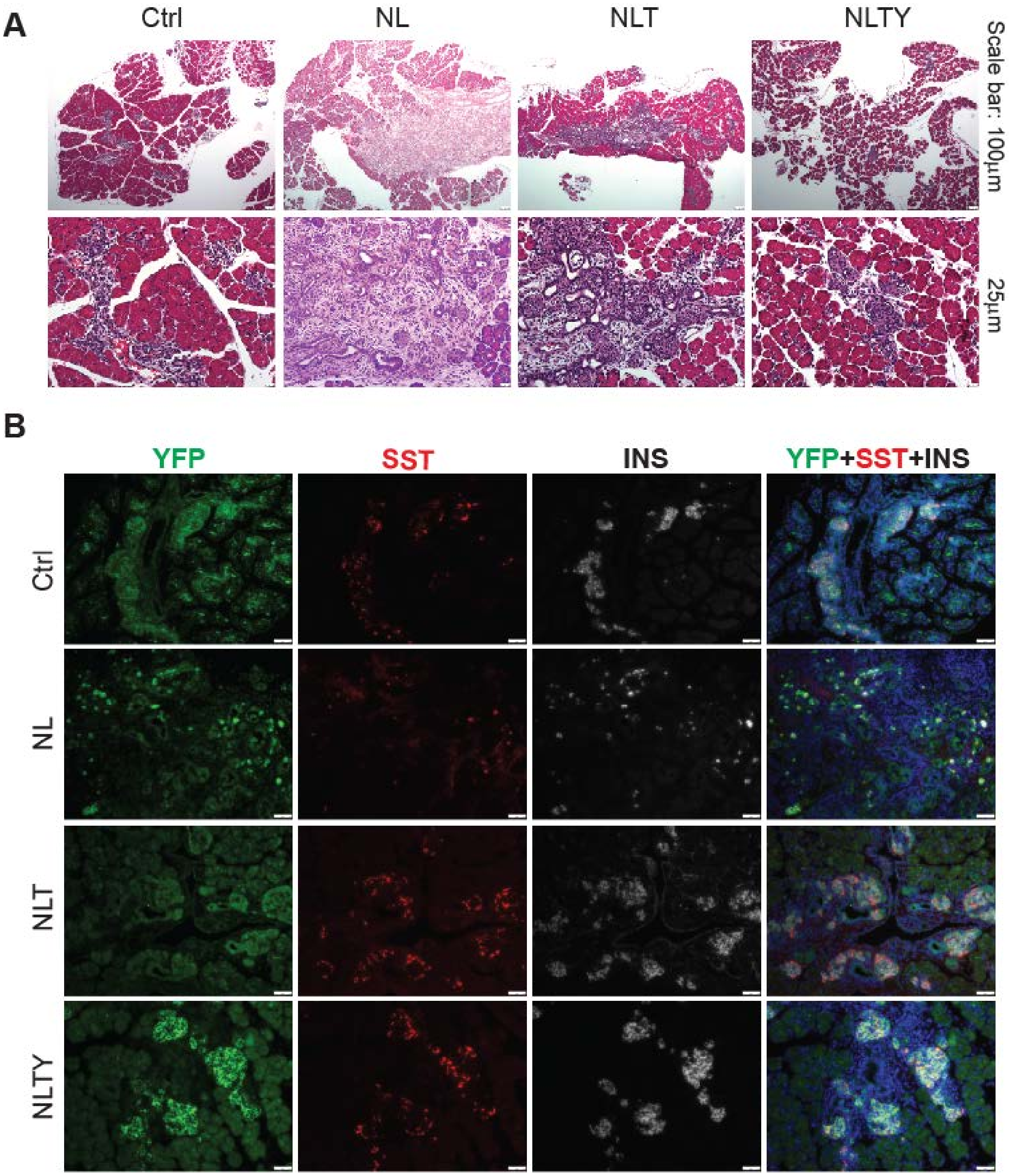


Figure S6. Removal of YAP1/TAZ rescued the defect in endocrine differentiation in NL pancreas. A. H&E staining showed morphologic defect in NLT pancreas, but appeared normal in NLTY pancreas. B. Co-staining of YFP, Insulin (INS), and Somatostatin (SST) showed that only a small fraction of YFP+ cells were also positive for INS or SST in NL pancreas, whereas most YFP+ cells in NLT and NLTY pancreas were INS or SST positive, more closely resembling the control phenotype.

Table S1. Primers for genotyping and qRT-PCR

| Gene name | Forward | Reverse |
|---|---|---|
| *Lats1* | GCGATGTCTAGCCCATTCTC | GGTTGTCCCACCAACATTTC |
| *Lats2* | AGCCTGACAACATACTCATCG | AATCCAGTGCAGAGGCCAAA |
| *Yap1* | TACTGATGCAGGTACTGCGG | TCAGGGATCTCAAAGGAGGAC |
| *TAZ* | GAAGGTGATGAATCAGCCTCTG | GTTCTGAGTCGGGTGGTTCTG |
| *Ankrd1* | TAATCGCTCACAATCTGTTGACA | GCCTCTCACCTTCCGACCT |
| *Ctgf* | GGCCTCTTCTGCGATTTCG | GCAGCTTGACCCTTCTCGG |
| *Cyr61* | TCCTCAGTGAGTTGCCCTC | CCCACCTAAGAGCCTCAGG |
| *Spp1* | AGCAAGAAACTCTTCCAAGCAA | GTGAGATTCGTCAGATTCATCCG |
| *Cxcl10* | CCAAGTGCTGCCGTCATTTTC | GGCTCGCAGGGATGATTTCAA |
| *Cxcl16* | CCTTGTCTCTTGCGTTCTTCC | TCCAAAGTACCCTGCGGTATC |
| *Tgfb1* | CCACCTGCAAGACCATCGAC | CTGGCGAGCCTTAGTTTGGAC |
| *Amy* | TTGCCAAGGAATGTGAGCGAT | CCAAGGTCTTGATGGGTTATGAA |
| *Ptf1a* | TCCCATCCCCTTACTTTGATGA | GTAGCAGTATTCGTGTAGCTGG |
| *Cpa1* | CAGTCTTCGGCAATGAGAACT | GGGAAGGGCACTCGAACATC |
| *Hnf1b* | AGGGAGGTGGTCGATGTCA | TCTGGACTGTCTGGTTGAACT |
| *Sox9* | AGTACCCGCATCTGCACAAC | ACGAAGGGTCTCTTCTCGCT |
| *Krt19* | GGGGGTTCAGTACGCATTGG | GAGGACGAGGTCACGAAGC |
| *Ins1* | CACTTCCTACCCCTGCTGG | ACCACAAAGATGCTGTTTGACA |
| *Ins2* | GCTTCTTCTACACACCCATGTC | AGCACTGATCTACAATGCCAC |
| *ChrA* | ATCCTCTCTATCCTGCGACAC | GGGCTCTGGTTCTCAAACACT |
| *GAPDH* | AGGTCGGTGTGAACGGATTTG | GGGGTCGTTGATGGCAACA |
